# Mirror-assisted light-sheet microscopy: a simple upgrade to enable bi-directional sample excitation

**DOI:** 10.1101/2024.04.05.588276

**Authors:** Asaph Zylbertal, Isaac H. Bianco

## Abstract

**Significance:** Light-sheet microscopy is a powerful imaging technique that achieves optical sectioning via selective illumination of individual sample planes. However, when the sample contains opaque or scattering tissue, the incident light-sheet may not be able to uniformly excite the entire sample. For example, in the context of larval zebrafish whole-brain imaging, occlusion by the eyes prevents access to a significant portion of the brain during common implementations using unidirectional illumination.

**Aim:** We introduce mirror-assisted light-sheet microscopy (mLSM), a simple inexpensive method that can be implemented on existing digitally scanned light-sheet setups by adding a single optical element – a mirrored micro-prism. The prism is placed near the sample to generate a second excitation path for accessing previously obstructed regions.

**Approach:** Scanning the laser beam onto the mirror generates a second, reflected illumination path that circumvents the occlusion. Online tuning of the scanning and laser power waveforms enables near uniform sample excitation with dual illumination.

**Results:** mLSM produces cellular-resolution images of zebrafish brain regions inaccessible to unidirectional illumination. Imaging quality in regions illuminated by the direct or reflected sheet is adjustable by translating the excitation objective. The prism does not interfere with visually guided behaviour.

**Conclusions:** mLSM presents an easy to implement, cost-effective way to upgrade an existing light-sheet system to obtain more imaging data from a biological sample.

## Introduction

Light-sheet fluorescence microscopy is a powerful imaging tool with a large variety of biosciences applications (Stelzer *et al*., 2021). It is based on exciting the sample with a thin light-sheet while imaging the emitted fluorescence via a perpendicular collection path. By restricting illumination to a single plane, light-sheet microscopes achieve optical sectioning while minimizing bleaching and photodamage. In a typical digitally scanned light-sheet setup, the sample is illuminated by a Gaussian beam that is rapidly scanned along the ‘y-axis’, perpendicular to beam direction, using a galvanometric mirror to excite a single sample plane. For volumetric imaging, multiple planes are exposed sequentially by displacing the light-sheet along the ‘z-axis’ using a second scan mirror.

A particularly successful application of light-sheet microscopy has been *in vivo* cellular-resolution functional imaging in the small, transparent brains of larval zebrafish expressing fluorescent calcium indicators (Ahrens *et al*., 2013), which has provided insights into the function of distributed neuronal circuits controlling vertebrate behaviour (Keller, Ahrens and Freeman, 2015; Dragomir, Štih and Portugues, 2020; Petrucco *et al*., 2023; Zylbertal and Bianco, 2023). In such experiments, the brain is illuminated from the side (laterally) and a light-sheet is formed by scanning the beam in a direction (‘y’) that corresponds to the rostrocaudal anatomical axis. However, in this arrangement, one of the eyes occludes the excitation beam and so a significant proportion of the total brain volume (∼20-25%, Ahrens *et al*., 2013) cannot be imaged.

Existing solutions to this problem include changing the geometry of the excitation path with respect to the animal and/or adding a second path. Specifically, in Dual-View Selective Plane Illumination Microscopy (diSPIM; Kumar *et al*., 2014) both the excitation and detection axes are coherently rotated by 45° and in Swept Confocally-Aligned Planar Excitation microscopy (SCAPE; Bouchard *et al*., 2015) a single objective positioned above the animal functions for fluorescence detection and sample excitation via an oblique plane. In other microscopes, a second excitation path has been constructed to illuminate the animal from an orthogonal “frontal” aspect, in addition to lateral illumination (Vladimirov *et al*., 2014). While these variants of light-sheet microscopy do improve optical access, those utilising oblique illumination require a completely new instrument design and cannot be easily implemented on existing set-ups. The addition of a second excitation path may be undesirable as it requires additional space for a second excitation objective (EO), which hinders presentation of visual stimuli in the animal’s frontal visual field. Such stimuli are commonly used, for example, in the context of virtual hunting assays (eg Bianco and Engert, 2015; Zylbertal and Bianco, 2023).

Here, we introduce mirror-assisted light-sheet microscopy (mLSM), as a simple and cost-effective solution that is easily implemented on existing light-sheet setups that employ unidirectional illumination. It is based on producing a second illumination path using a mirrored micro-prism, with corresponding adjustments to beam scanning and laser power modulation (Figure 1B). We demonstrate that this enables imaging of regions normally obscured during unidirectional illumination. In addition, we show that image quality may be adjusted by displacing the EO and comparable, near-cellular resolution data can be obtained from tissue that is illuminated directly, or via the reflected sheet. Finally, we show that the micro-prism does not interfere with larval zebrafish responding to stimuli in their frontal visual field.

**Figure 1:**
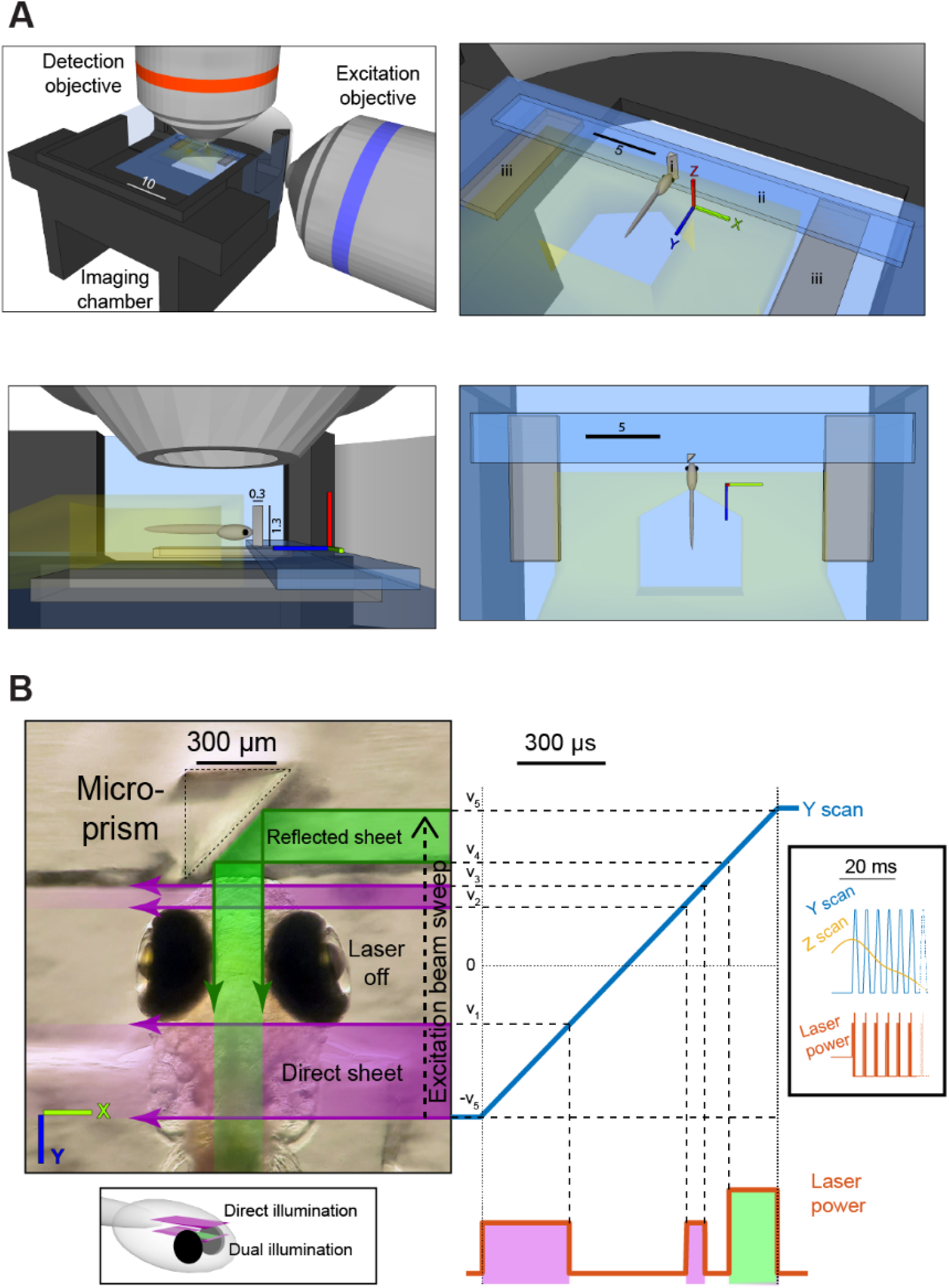
Overview of mLSM. (**A**) Schematic of the imaging chamber and the orthogonal excitation and detection objectives (top-left); isometric (top-right), side (bottom-left) and top (bottom-right) views of the fish and prism (i), glass strip (ii) and plastic bases (iii). Glass is depicted in blue and agarose in yellow, distances indicated in millimetres. (**B**) Left - Zebrafish larva and micro-prism, overlayed with excitation laser paths for exposure of a dual illumination plane. Right - analogue voltage waveforms for y scan mirror (blue) and laser power (orange) for imaging of a dual illumination plane. Magenta and green shading indicate direct (lateral) and reflected (frontal) sheets, respectively. v_1-5_ – y mirror voltages corresponding to the posterior and anterior edges of the eyes (v_1/2_), the rostral edge of the brain (v_3_), and the medial surfaces of the eyes (v_4/5_). Bottom inset: diagrammatic location of representative direct and dual illuminated planes. Right inset: Analog voltage waveforms for a volume scan, including multiple scan planes and the z mirror waveform (yellow).

## Materials and Methods

### Animals

Zebrafish (*Danio rerio*) larvae were reared on a 14/10 hr light/dark cycle at 28.5 °C. For all experiments, we used zebrafish larvae homozygous for the *mitfa*^*w2*^ skin-pigmentation mutation (Lister *et al*., 1999) and *Tg(elavI3:H2B-GCaMP6s)*^*jf5Tg*^ transgene (Vladimirov *et al*., 2014, ZFIN ID: ZDB-ALT-141023–2). All larvae were fed *Paramecia* from 4 dpf onward. Animal handling and experimental procedures were approved by the UCL Animal Welfare Ethical Review Body and the UK Home Office under the Animal (Scientific Procedures) Act 1986.

### Light-sheet microscopy setup

We implemented mLSM on a custom-built digitally scanned light-sheet microscope that was in routine use in the lab (Zylbertal and Bianco, 2023). The excitation path included a 488 nm laser source with an analogue input for power modulation (OBIS 488-50 LX, Coherent, Santa Clara, California), a pair of galvanometer scan mirrors (6210H, Cambridge Technology, Bedford, Massachusetts) and objective (Plan 4X, 4x/0.1 NA, Olympus, Tokyo, Japan), with its back aperture limited to 5.5 mm using an adjustable iris (for an effective NA of ∼0.06). One mirror was used to scan the beam along the y-axis to produce a light-sheet during a single camera exposure and the second mirror displaced the light-sheet along the z-axis across frames. Zebrafish larvae were mounted such that the y-axis corresponds to the anatomical rostrocaudal axis and the zaxis to the dorsoventral axis.

The orthogonal detection path, parallel to the ‘z’ axis, comprised a water-immersion detection objective (XLUMPLFLN, 20x/1.0 NA, Olympus), a telecentric tube lens (TTL200MP, Thorlabs, Newton, New Jersey, f=200 mm), two relay lenses (a 2X apochromatic objective, TL2X-SAP, Thorlabs, f=100 mm, and a 50 mm Ø achromatic doublet, G322303000, Excelitas Technologies, Waltham, Massachusetts, f=120 mm) in a 4f configuration, and sCMOS camera (Kinetix, Teledyne Photometrics, Tucson, Arizona). For remote focusing (Fahrbach et al., 2013), an electrically tuneable lens (ETL, EL-16-40-TC-VIS-20D, Optotune, Dietikon, Switzerland) was installed between the relay lenses, conjugate to the back focal plane of the detection objective.

### Prism assembly preparation

The core component enabling mLSM is a custom fabricated 0.3x0.3x1.3 mm right-angle prism (45°-45°-90°), with its hypotenuse surface coated with aluminium for external reflection (Figure 1A, i, Precision Optics Corporation, Gardner, Massachusetts). To prepare it for use in imaging experiments, we first glued the prism upright on a 3x18 mm strip of glass (cut from a 18x18 mm microscope coverslip), using UV-cured optical adhesive (#61, Noroland, Jamesburg, New Jersey, Figure 1A, ii). The glass strip was glued to two 2x5x0.75 mm clear PETG plastic sheets, that function as bases for the prism assembly (Figure 1A, iii). Each prism assembly was then used repeatedly in multiple imaging sessions.

### Animal mounting and prism positioning

Larval zebrafish were mounted at 5 dpf in 3% low melting point agarose (Sigma-Aldrich, St. Louis, Missouri) on top of a 22x22 mm coverslip. They were allowed to recover overnight before imaging at 6 dpf. Prior to the imaging session, the prism assembly was attached to the mounting coverslip using a small amount of silicone grease that was applied to its plastic bases. The coverslip with the prism assembly was then moved to a custom 3D printed chamber (SLS Nylon 12, 3DPRINTUK, London, United Kingdom) and the prism assembly position was fine-tuned to make sure the prism is oriented correctly and placed as close as possible to the fish but without touching it (Figure 1A-B). The long dimension of the prism (1.3mm, its ‘height’, along the z axis) enabled flexibility in the mounting depth of the fish.

To fine-tune prism position prior to imaging, a DC bias voltage was sent to the ‘y’ scan mirror to displace the beam onto the prism, enabling inspection of its reflected portion (using diluted fluoresceine placed in the bath). Small errors in the angle of the prism with respect to the base coverslip would often cause the reflected beam to slant with respect to the imaging plane. We diagnosed beam slanting by translating the beam waist (by moving the EO along the optical axis) and re-focusing the detection objective and corrected it by pitching the imaging chamber slightly using coverslips stacked beneath it (alternatively, this may be fixed by rotating the stage about the x axis).

### Image Acquisition

All microscope control was implemented using custom LabVIEW (National Instruments, Austin, Texas) routines.

To assess image quality and make point-spread function measurements, we acquired volumetric image stacks of brains or fluorescent beads using stepwise change in z-mirror position and corresponding ETL optical power. Brain anatomical stacks were comprised of 185 planes spaced 0.93 μm apart, each obtained by averaging 20 images (130 μW laser power at sample, 1 ms exposure). Image stacks of fluorescent beads were comprised of 200 planes spaced 0.2 μm apart, each obtained by averaging 25 images (520 μW laser power at sample, 35 ms exposure).

For functional calcium imaging, we acquired image volumes by dynamically changing ETL power and z-mirror position (see waveform in Figure 1B), at 5 volumes per second. Each volume was comprised of 37 imaging planes spaced 3.6 μm apart. Each plane was exposed to 130 μW laser power for 1 ms.

### Scan waveform adjustment

For dual illumination via direct and reflected light-sheets, we adapted the voltage waveforms sent to the scan mirrors and the analogue laser power control signal (Figure 1B). To accommodate variation in the precise geometry of the fish with respect to the prism across sessions, the software enabled fine-tuning of the waveforms properties, with online feedback. First, while viewing an imaging plane dorsal to the eyes, we tuned the y scan amplitude (Figure 1B, +/-v_5_), ensuring the reflected sheet reached the medial surface of the eye proximal to the EO. Next, we tuned the voltages corresponding to the rostral edge of the brain (v_3_), the side of the reflected sheet distal to the EO (v_4_), and the approximate posterior and anterior edges of the eyes (v_1/2_). Lastly, while viewing a plane close to the dorsal edge of the eyes, we tuned the z scan voltage corresponding to the dorsal edge of the eyes, so as to divide the imaging planes into ‘direct illumination only’ planes and ‘dual illumination’ planes.

For direct illumination planes, the laser was on during the entire sweep across the brain (-v_5_ to v_3_). For dual illumination planes, the laser power was modulated to generate the reflected sheet (laser on between v_4_ and v_5_) and blanked while scanning over the eye (v_1_-v_2_). The laser power was also increased between v_4_ and v_5_ to compensate for the power lost in reflection.

### Point Spread Function (PSF) measurement

We used 0.1 μm blue/green/orange/dark red fluorescent beads (T7284, Invitrogen, Waltham, Massachusetts) embedded in a block of 1% low melting point agarose (2.5 μl bead stock in 500 ml agarose). The spacing between prism and agarose was adjusted to correspond to the rostral part of the brain and 250x125x50 μm stepwise stacks were acquired (1024x512 pixels lateral field of view, 250 axial steps of 0.2 μm), while the sample was illuminated by either the reflected or direct sheet for 4 different EO positions. For each illumination direction and objective distance, we chose 35 beads for PSF analysis. To restrict the analysis to beads illuminated by an approximately uniform-width light-sheet, all chosen beads were within 18 μm from the beam waist along the beam propagation axis (ie a small fraction of the 118 μm Rayleigh length). We then calculated the full width at half maximum (FWHM) from gaussian fits to the axial intensity profile around each bead. To measure the axial PSF of the detection optics alone, we acquired stacks by stepping the ETL optical power without changing the z position of the light-sheet.

### Assessment of biological images

To quantify image quality with emphasis on biologically relevant spatial scales, we combined two approaches: The Useful Contrast (UC) metric proposed by Truong *et al*. (2011), and automatic anatomy-based segmentation of images to detect regions of interest (ROIs) corresponding to neuronal nuclei (Figure 2, Kawashima *et al*., 2016).

**Figure 2:**
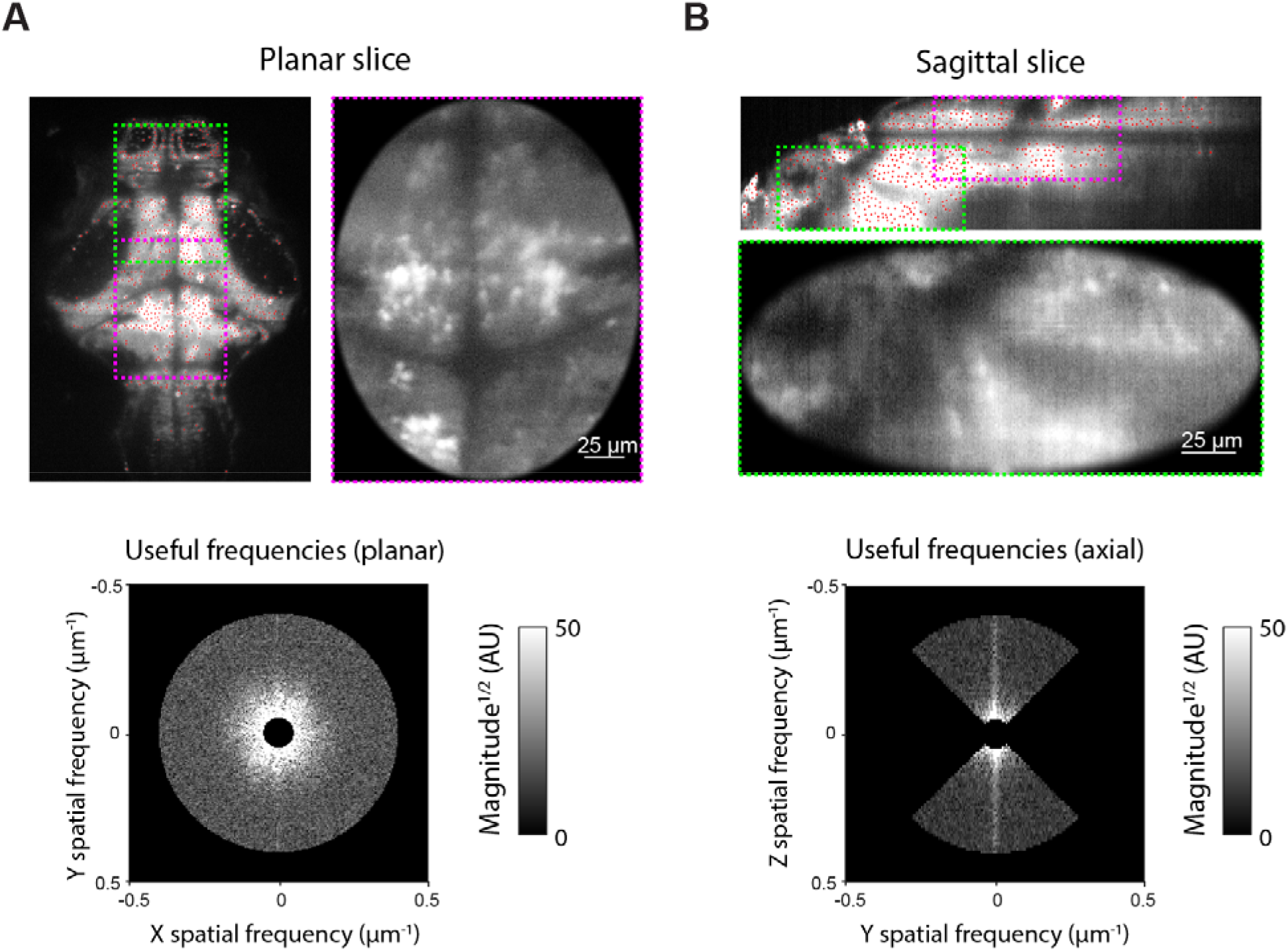
Image quality quantification. (**A**) Left: Raw image plane with detected ROIs indicated by red dots. Regions used for UC metric calculation are marked (direct illumination – magenta; reflected illumination – green). Right: Cropped image region following ‘soft’ brightness truncation and windowing. Bottom: Power spectrum for the cropped image, showing biologically relevant frequencies used to calculate the planar UC. (**B**) Same as (A), for a reconstructed sagittal slice. Only spatial frequencies in the axial (Z) direction were taken for axial UC calculation.

For UC metric calculation, we first applied a ‘soft’ truncation of pixel grayscale values to avoid bias by localised features with extreme brightness values. The pixel grey values ‘X’ were transformed by 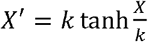, with the maximal brightness k chosen to be the top 0.1 brightness percentile (across the volume). Next, 200x240x100 μm sub-volumes were selected from the medial dorsocaudal or ventrorostral part of the brain (to assess direct and reflected illumination, respectively, Figure 2A-B). Individual planes were windowed with a soft-edged oval mask to avoid edge effects. To calculate planar UC, the 2D power spectrum of each plane was calculated, and the power at biologically relevant spatial frequencies (0.05 – 0.4 μm^-1^, corresponding to a spatial scale of 2.5 – 20 μm) was normalised by the total power. Axial UC was obtained similarly by using sagittal or coronal slices reconstructed as a cross-section along imaging planes and taking the power at the axial direction alone (Figure 2B). To provide feedback during EO position adjustment, planar UC was calculated online by the microscope control software during rapid automated switching between direct and reflected illumination of a single plane.

To detect neuronal nuclei in images, every 20^th^ image plane was automatically segmented using the algorithm proposed by Kawashima *et al*. (2016). This algorithm assumes individual neurons appear as discs with radius r (we used r=2.5 μm to match the average neuron size). Briefly, it first extracts regions in the image with high local contrast, and extracted pixels are then intensity-normalised based on a surrounding image patch (a circular region with a radius of 4r/3). The normalized image is then further smoothed by circular kernel (with a radius of 0.5r) and ROI centroids are defined as locations of peak brightness within discs of radius r (red dots in Figure 2). All image analysis was done in Matlab.

### Visual stimulus presentation and response analysis

Visual stimuli were back-projected (DLP LightCrafter E4500 MKII 385 nm UV, EKB Technologies Ltd, Bat Yam, Israel) onto a curved screen forming the wall of the imaging chamber in front of the animal, at a viewing distance of 18 mm. Visual stimuli were designed in Matlab using Psychophysics toolbox (Brainard, 1997). Stimuli comprised 4° UV bright spots on a red background, moving along a circular trajectory (r = 3°) at 5 rotations/s for 2 seconds, centred at an azimuth position of -40°, -25°, -10°, 0°, 10°, 25° or 40° (pseudo-random order, 5 repetitions for each direction in total).

Eye position was tracked at 50 Hz under 850 nm illumination using a sub-stage GS3-U3-41C6NIR-C camera (Point Grey, Richmond, Canada). Eye movements were categorized as a convergent saccade if both eyes made nasally directed saccades within 150 ms of one another (Dowell, Lau and Bianco, 2023). Stimulus presentation and behaviour tracking were implemented using LabVIEW and Matlab.

For details on functional calcium imaging analysis and image registration (Figure 6C) see Zylbertal and Bianco (2023).

## Results

When applying mLSM, image volumes can be assembled using two types of imaging planes. For imaging the brain of larval zebrafish, dorsal planes, located above the eyes, are imaged using direct (lateral) illumination, with constant laser power during the y-mirror scan, as in conventional light-sheet imaging. However, ventral planes, which would ordinarily be occluded by the eye, are imaged with dual illumination: Each exposure combines direct (lateral) as well as reflected (frontal) light-sheet excitation during a single camera frame (Figure 1B).

Inspection of raw image planes acquired with dual illumination (Figure 3, Videos 1-2) showed that anatomical features could be clearly resolved in the portion of the zebrafish brain that was exclusively illuminated by the reflected sheet. This region comprises 25% of the imaged brain volume (1x10^7^ out of 3.9x10^7^ μm^3^) and includes 20% of all automatically-segmented ROIs (1.4x10^4^ out of 7x10^4^ ROIs, corresponding to neuronal nuclei).

**Figure 3:**
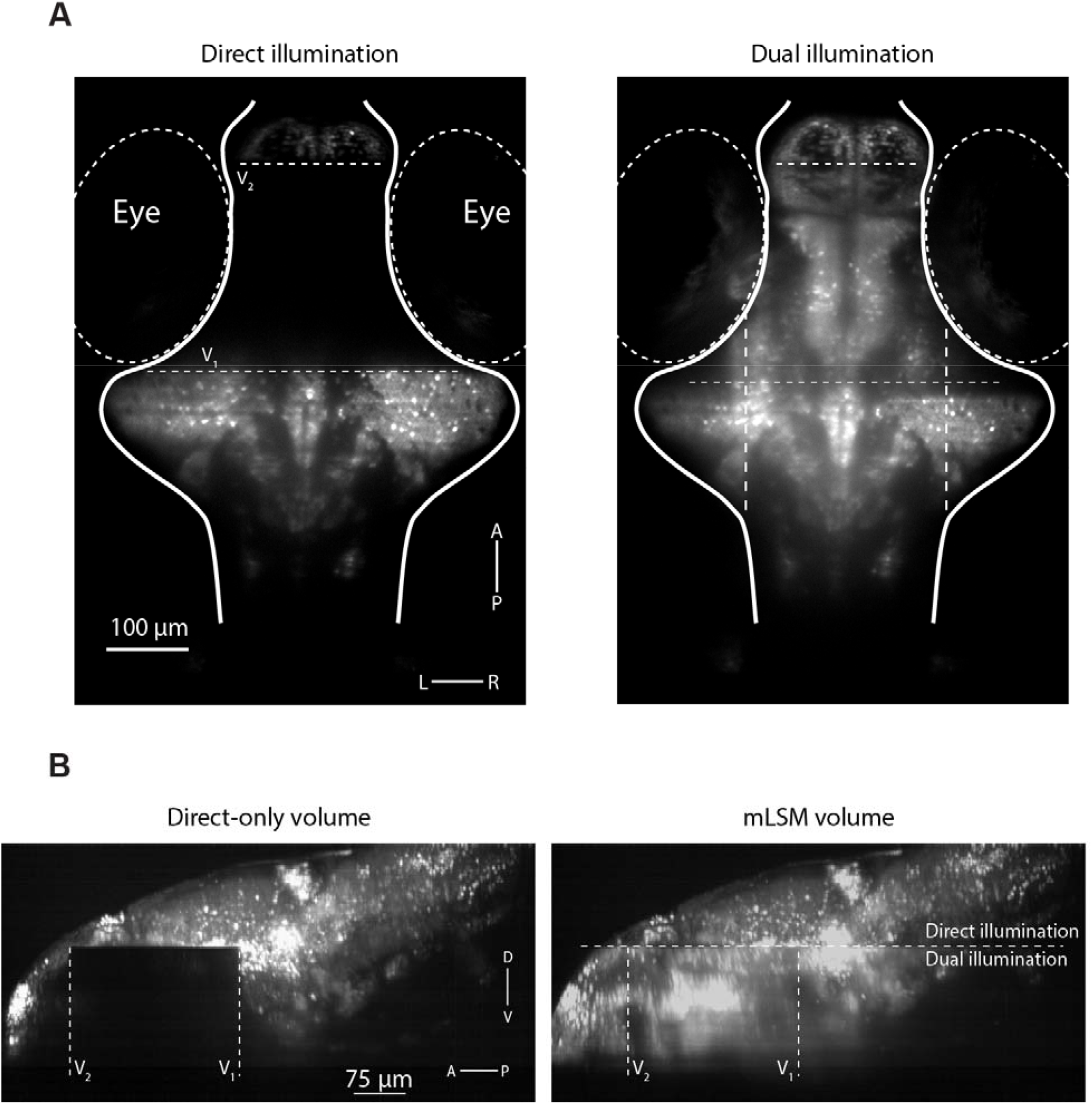
Mirror-assisted light-sheet imaging. (**A**) A single ventral imaging plane, acquired with conventional, direct illumination, in which the laser is off while scanning over the eyes (left) and with mLSM (dual illumination, right). (**B**) Sagittal maximal-intensity projection of a whole-brain volume acquired with direct-only (conventional) illumination (left) and the same volume acquired with mLSM (green). **Video 1:** Stacks acquired with dynamic Z scanning with direct-only illumination (left) and mLSM (right). **Video 2:** 3D maximal-intensity projection of a whole-brain volume acquired with direct-only illumination (top) and the same volume acquired with mLSM (bottom).

### Image quality considerations

For mLSM, both direct and reflected light-sheets derive from the same EO and, in our implementation, illuminate the specimen during a single camera exposure. Both features have implications for image quality that we consider in this section.

Because a single EO is used to generate both the direct and reflected light-sheets, a limitation of mLSM is that the Gaussian beam profile cannot be independently adjusted for the two excitation paths. Specifically, the gaussian beam width, w, reaches a minimal value (w_o_) at the focal plane of the EO, and the field of view (FOV) can be defined as the beam propagation distance over which the sheet is less than twice this minimal waist thickness (Figure 4A). By adjusting the excitation NA (using an iris) a compromise between w_0_ and FOV must be found such that a sufficiently thin light sheet (of similar width to the biological features of interest) extends over a FOV that is adequate to cover the specimen of interest. Thus, a higher NA reduces w_0_ but also the FOV, whereas a lower NA increases both. In our microscope, we used a 5.0 mm iris to set the excitation NA to approximately 0.06. For the resulting Gaussian beam we measured w_o_ = 4.7 μm (full width at half max, FWHM) and FOV = 346 μm, similar to theoretical values of 2.7 μm and 350 μm respectively (for 488 nm excitation wavelength). With this beam geometry and the EO at its native position (i.e. where the beam waist is centred at the midline of the brain), the direct illumination path produced a light-sheet that was sufficiently thin to image zebrafish neurons across the width of the brain. However, in the reflected beam path, the additional prism-to-specimen distance (∼120 μm) means the tissue is illuminated by a light-sheet that is thicker than 2·w_0_ (Figure 4B). A thinner reflected light-sheet can be obtained by translating EO towards the specimen, but this will shift the beam waist away from the centre of the specimen in the direct illumination path.

**Figure 4:**
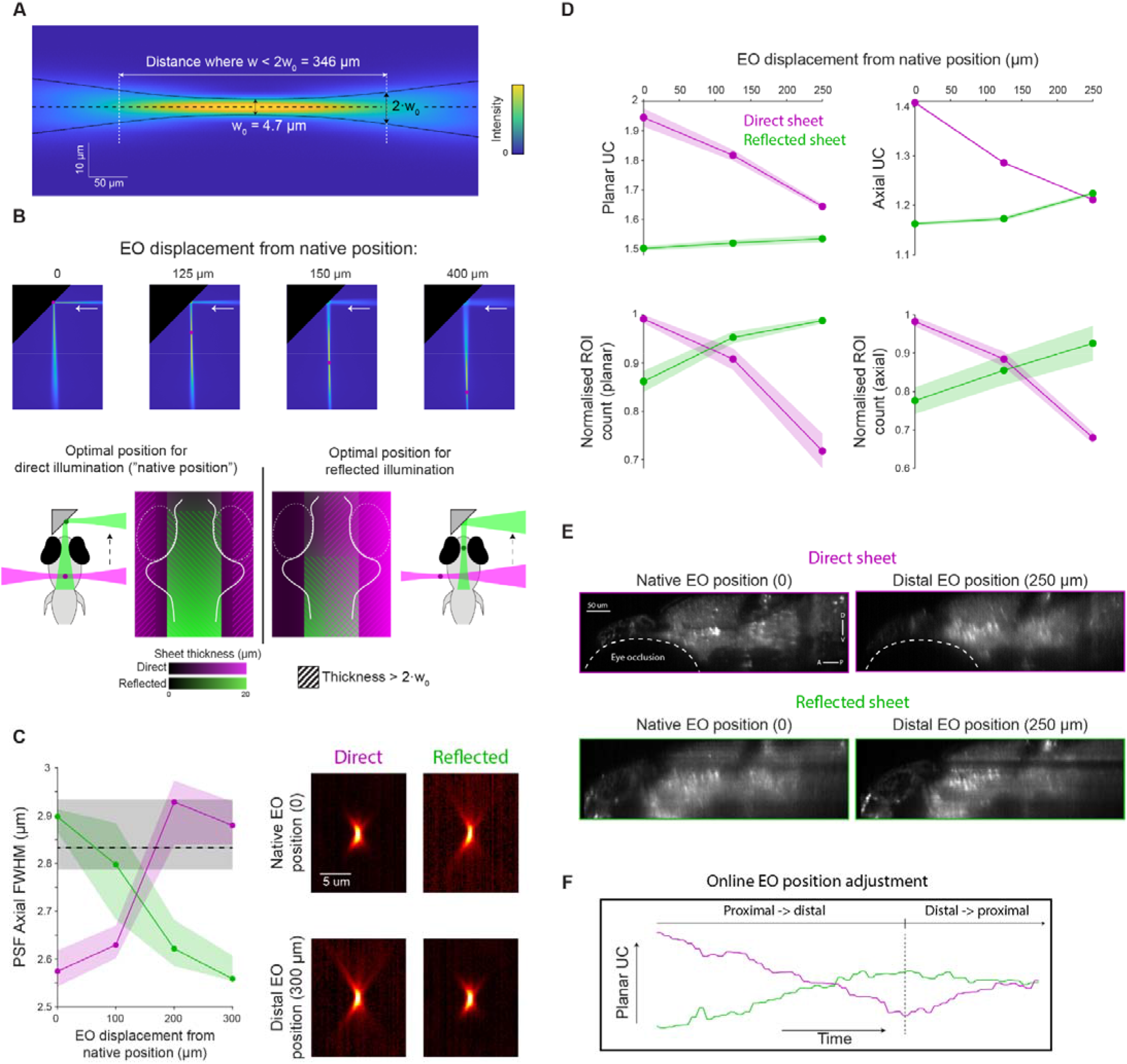
Image quality is sensitive to the geometry and position of direct and reflected beams. (**A**) Measured gaussian beam properties (direct illumination of fluoresceine). (**B**) Translating the EO towards the specimen repositions the waist of the direct and reflected beams. Top: views of the reflected beam at four different EO positions (red circles denote the beam waist). At the native position (“zero”), the waist of the direct beam is centred at the midline of the brain. Bottom: The trade-off between direct and reflected sheet thickness, at different EO positions. (**C**) Left: PSF axial size as a function of EO position with direct (magenta) or reflected (green) illumination (N=35 beads). Black dashed line indicates the axial PSF of the detection optics alone. Lines show median with 40^th^-60^th^ percentiles shaded. Right: PSF X-Z profile acquired with direct (left) and reflected (right) illumination, with a native (top) and distal (bottom) EO positions (median images, gamma-corrected, γ=0.7). (**D**) Image quality as a function of EO position, as measured by UC (top), or number of detected ROIs (bottom, normalised by the maximal number of ROIs detected for each illumination direction). Left: Planar quality measures, averaged across planes. Right: Axial quality measures, averaged across sagittal slices. Shaded area indicates SEM across slices. (**E**) Example sagittal section acquired with direct (top) and reflected (bottom) illumination, at native (left) and distal (right) EO positions. (**F**) Example of online assessment of UC while manually adjusting EO position to achieve approximately equal image quality for direct and reflected illumination. A, Anterior; P, Posterior; D, Dorsal; V, Ventral.

To evaluate this trade-off, we first made point-spread function measurements to empirically verify the axial resolution obtained with direct and reflected illumination. We obtained image stacks of a field of fluorescent beads, under either direct or reflected illumination, at four different positions of the EO. As expected, (1) the axial FWHM increased for beads illuminated by thicker parts of either beam (i.e. further away from the beam waist, w_0_) and (2) translating the EO towards the sample improved axial resolution for the reflected light-sheet but degraded it for the direct light-sheet. We note that the axial PSF is bounded by the high-NA detection objective (NA = 1.0, ∼2.8 μm; dashed line in figure 4C) such that near-cellular resolution imaging is still obtained in regions of the sample far from the beam waist, using either direct or reflected illumination.

When imaging biological samples, excitation light is scattered as it propagates through tissue such that real imaging performance invariably deviates from ideal conditions. We therefore used two metrics to quantitatively assess the quality of biological images obtained with each excitation path. First, we applied the Useful Contrast (UC) metric proposed by Truong *et al*. (2011) which measures the spatial frequency content of images at biologically relevant length scales, which is a function of resolution, contrast, and signal-to-noise ratio (Materials and Methods). Second, we tested how imaging conditions affect the automatic detection of putative neuronal ROIs using a contrast-based segmentation algorithm (Kawashima *et al*., 2016). We acquired image volumes spanning the entire larval zebrafish brain with either direct or reflected illumination, at three different EO positions and calculated the planar UC (averaged across planes, Figure 2A), the axial UC (averaged across sagittal slices, Figure 2B) and applied the segmentation algorithm on every 20^th^ plane or sagittal slice. In agreement with theory and measured changes in PSF, both metrics showed that translating the EO away from its ‘native position’ improves image quality under reflected illumination at the expense of image quality under direct illumination (Figure 4D-E). However, in general, image quality under reflected illumination was somewhat less sensitive to EO position, which might be because it is bounded by the scattering properties of anterior tissue along the path of the reflected illumination (e.g. the animal’s jaw). This is in agreement with previous reports, where lateral illumination of the larval zebrafish brain generated a greater proportion of cellular resolution data (92% of the imaged volume; Ahrens *et al*., 2013) as compared with anterior illumination (80%; Vladimirov *et al*., 2014).

In sum, in mLSM a balance must be struck between image quality obtained using direct versus reflected light-sheets, which is controlled by adjusting EO position. For specific research applications, this balance will depend on factors including what fraction of the sample can be accessed by each light-sheet and in what regions of the sample the highest image quality is required. We also note that the relative position of the prism with respect to the sample can vary slightly from one experiment to the next. For these reasons, we implemented online UC calculation within our microscope control software (Materials and Methods), to provide the experimenter with immediate feedback during manual translation of the EO such that image quality can be easily tuned for each and every experiment.

In our implementation of mLSM, we chose to combine direct and reflected light-sheet illumination within individual camera frames, in order to maximise frame rate for functional imaging. A consequence of this is that some regions of the sample are illuminated bi-directionally and in those regions, excitation by the thicker light-sheet is expected to degrade axial resolution. To assess this issue under routine imaging conditions, we compared UC for direct-only versus dual illumination in the region of the sample that is excited by both beam paths (yellow box in Figure 5A). Our results show that the addition of the reflected sheet had only a modest effect on axial UC and it was restricted to around 50 μm at the rostral edge of the overlap region. We conclude that, at least for this application, the use of dual illumination has only a modest impact on image quality while providing access to a substantial portion of the brain that is ordinarily inaccessible.

**Figure 5:**
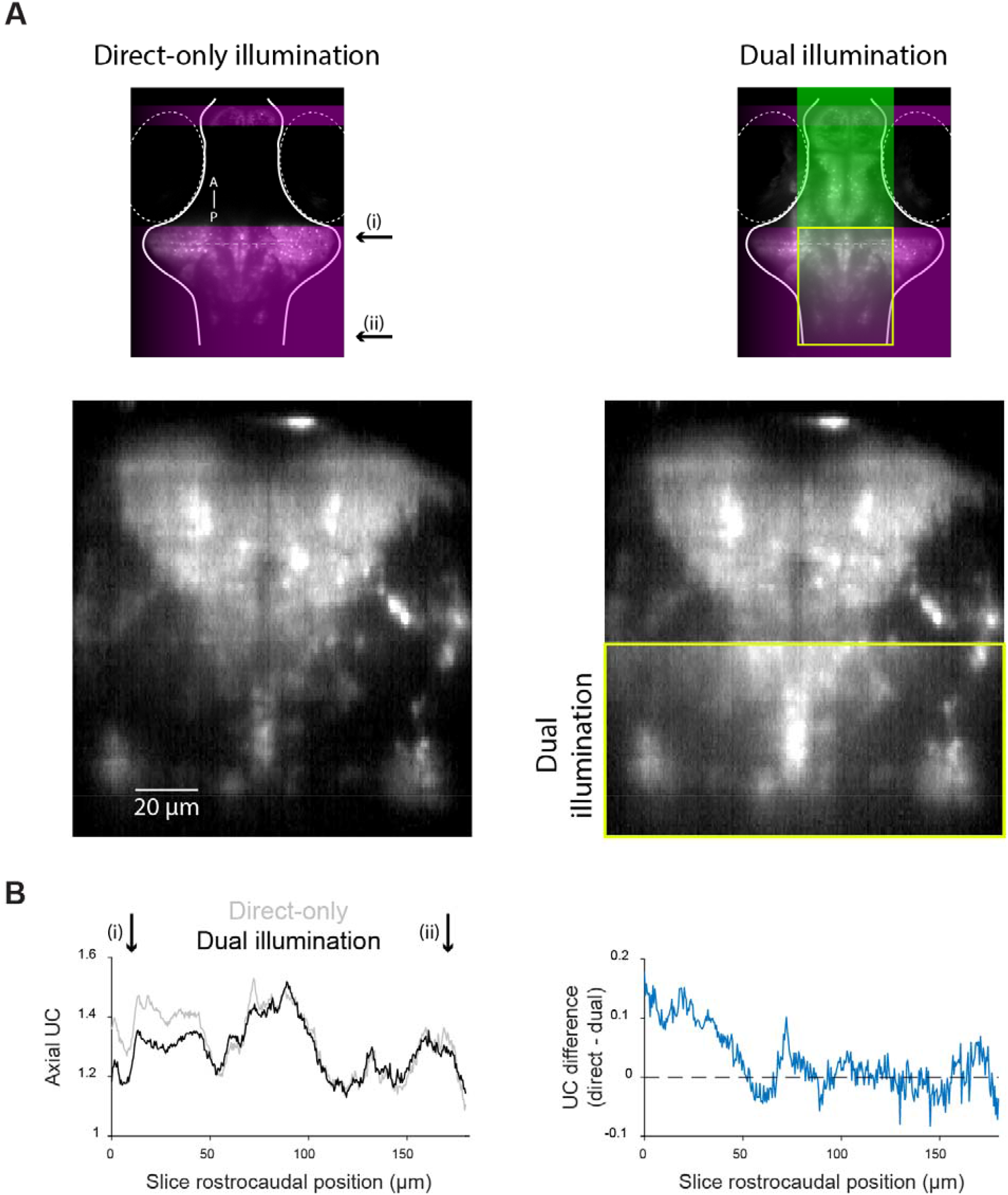
The effect of dual illumination on image quality. (**A**) Top: a ventral imaging plane with direct-only (left) or dual illumination (right). Bottom: A coronal slice at the level indicated by dashed line in top panels. Yellow box indicates the region of overlapping illumination. (**B**) Left: Axial UC as a function of rostrocaudal position (from i—ii, indicated in A) within the overlap region. Right: Difference in UC between dual illumination and direct-only illumination.

### Visuomotor behaviour

A potential concern with mLSM is that although the micro-prism is very small, by placing it in front of the fish it may nonetheless obstruct the animal’s visual field, in turn interfering with analysis of visually evoked neural activity and behaviour. We calculated that with typical prism positioning and mean ocular vergence angle (21.1°, Bianco, Kampff and Engert, 2011), the prism should be just outside the visual field and therefore have only minimal effects (Figure 6A). To test this, we presented 10 larval zebrafish with prey-like moving spots centred at seven different positions across the animals’ frontal visual field (ranging from -40° to +40° azimuth) while tracking their behaviour and performing imaging to assess visually evoked neural activity. The protocol was repeated twice for each fish, in the presence and absence of the micro-prism.

**Figure 6:**
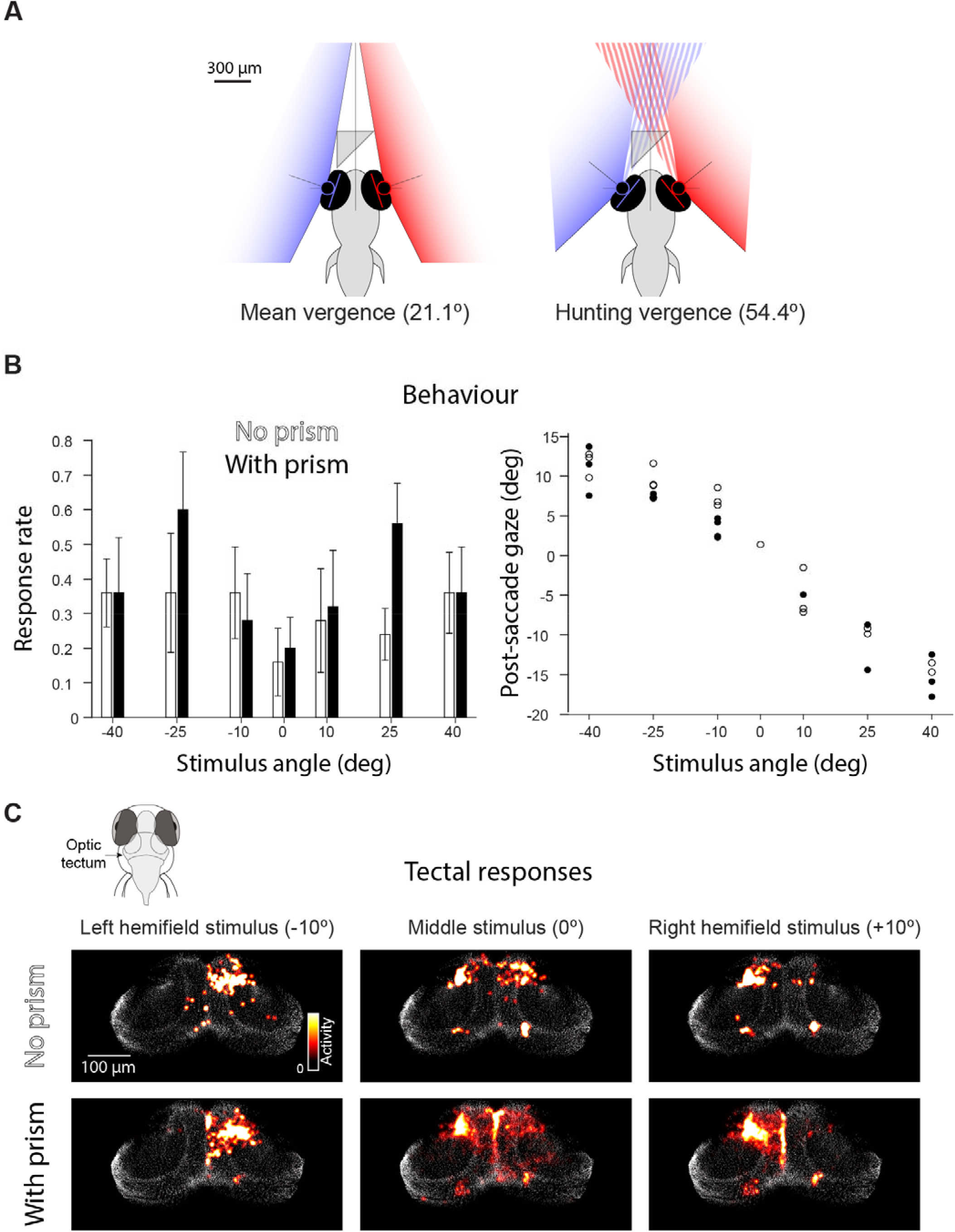
The micro-prism does not obstruct or distort the visual scene. (**A**) Predicted impact of the micro-prism on the visual field during normal vergence (left) and following a convergent saccade (right) (Bianco, Kampff and Engert, 2011). (**B**) Left: Rate of hunting (convergent saccade) responses to rotating spot stimuli presented at different azimuth angles in the absence (white) and presence (black) of the prism (N=5 fish, error bars indicate SEM). Right: Post-saccade gaze angles in a single fish in the absence (empty circles) and presence (filled circles) of the micro-prism. (**C**) Neural activity in the optic tectum evoked by presentation of -10°, 0° and +10° stimuli for one fish in the absence (top) and presence (bottom) of the micro-prism. Each panel shows activity averaged over the 2 s of stimulus presentation (median of 5 repetitions).

The presence of the prism did not affect behaviour or visually evoked neural activity. Specifically, the probability of visually evoked hunting responses, as identified by convergent saccades, as well as their directionality were comparable in the presence versus the absence of the microprism (Figure 6B). Furthermore, we observed similar visually evoked neural activity in the optic tectum for stimuli presented across the range of azimuth positions (Figure 6C). The prism is unlikely to influence the lower and lateral visual field and accordingly we did not observe a difference in the optomotor responses evoked by drifting gratings presented below the animal (data not shown). Taken together, these observations suggest the prism does not significantly obstruct or distort the visual scene. However, we note that when fish initiate hunting, eye convergence substantially increase the size and proximity of the binocular visual field (Bianco, Kampff and Engert, 2011; Dowell, Lau and Bianco, 2023). Under these conditions, we estimate the prism occludes ∼20° of the visual field (Figure 6A), potentially impacting experiments involving hunting sequences (Discussion).

## Discussion

In this study, we demonstrated that a simple adaptation to a digitally scanned light-sheet microscope enables cellular-resolution imaging in regions of a sample that are normally occluded during unidirectional illumination. By positioning a mirrored micro-prism in front of a larval zebrafish and adapting the scanning waveforms accordingly, we were able to produce a second reflected light-sheet and gain access to the 20-25% of brain tissue that lies between the eyes. Importantly, this simple and inexpensive upgrade can be readily implemented in existing setups, including simple open-source designs (Gualda *et al*., 2013; Pitrone *et al*., 2013), by adding a single element. By avoiding the need for a second excitation objective, visual stimuli may be readily presented on a screen in front of the animal. Although we developed this approach for functional calcium imaging in larval zebrafish, it could be used in other cases where a second excitation path would provide access to a greater portion of the sample tissue.

In our implementation, the different path lengths for the direct and reflected beam resulted in a trade-off in image quality obtained using the two light-sheets. However, when imaging real biological samples we found that mLSM was still able to produce near-cellular resolution images including in regions of the sample illuminated exclusively by the reflected sheet, or illuminated by both. In future implementations, the excitation path could be enhanced, for example by the addition of an ETL or piezoelectric actuator, to rapidly shift the beam waist between the direct and reflected sheets. This would require a sufficiently slow frame scan, or acquisition of separate image frames for direct vs reflected illumination. This latter approach could also be used to prevent degradation of axial resolution in regions of the sample illuminated by both sheets, although we found that in practise this effect was minor and we opted not to reduce acquisition frame rates.

Returning to the use case of imaging neural activity during visuomotor behaviour in larval zebrafish, geometrical considerations indicated that the tiny micro-prism should minimally interfere with the animals’ frontal visual field. This was confirmed by observing visually evoked behaviour and tectal activity (Figure 6). However, an occlusion of at least 20° from the nasal side of the visual field of each eye is likely during hunting sequences where the eyes are converged (Bianco, Kampff and Engert, 2011; Patterson *et al*., 2013). A potential workaround would be to implement mLSM using a smaller prism. A 150 μm-edged prism, for example, would still produce a sufficiently wide reflected sheet and greatly reduce occlusion of the visual field, even when the eyes are converged.

In conclusion, mLSM is a simple, versatile, and robust method for adding a second illumination axis to existing digitally scanned light-sheet microscopes. It is suited for cases where opaque tissue prevents uniform illumination of the sample and can be used in conjunction with visual stimulus delivery and behavioural assays in larval zebrafish.

## Disclosures

The authors declare that no competing interests exist.

## Acknowledgements

The authors thank the members of the Bianco lab for helpful discussions and critical feedback on the manuscript and UCL Fish Facility staff for fish care and husbandry. This research was funded in whole, or in part, by the Wellcome Trust [220273/Z/20/Z]. For the purpose of Open Access, the author has applied a CC BY public copyright licence to any Author Accepted Manuscript version arising from this submission. This work was also supported by a Leverhulme Trust Project Grant [RPG-2023-041] and a BBSRC Discovery Fellowship [BB/S010564/1].

## Notes

### Competing Interest Statement

The authors have declared no competing interest.

### Summary of Updates

Revised fonts to improve PDF appearance (no content change)

## References

Ahrens, M. B. et al. (2013) “Whole-brain functional imaging at cellular resolution using light-sheet microscopy,” Nature methods, 10(5), pp. 413–420.

Antinucci, P., Folgueira, M. and Bianco, I. H. (2019) “Pretectal neurons control hunting behaviour,” eLife. eLife Sciences Publications Ltd, 8. doi: 10.7554/eLife.48114.

Bianco, I. H. and Engert, F. (2015) “Visuomotor transformations underlying hunting behavior in zebrafish,” Current biology: CB, 25(7), pp. 831–846.

Bianco, I. H., Kampff, A. R. and Engert, F. (2011) “Prey Capture Behavior Evoked by Simple Visual Stimuli in Larval Zebrafish,” Frontiers in systems neuroscience. Frontiers, 5, p. 101.

Bouchard, M. B. et al. (2015) “Swept confocally-aligned planar excitation (SCAPE) microscopy for high speed volumetric imaging of behaving organisms,” Nature photonics, 9(2), pp. 113–119.

Dowell, C. K., Lau, J. Y. N. and Bianco, I. H. (2023) “The saccadic repertoire of larval zebrafish reveals kinematically distinct saccades that are used in specific behavioural contexts,” bioRxiv. doi: 10.1101/2023.11.07.565345.

Dragomir, E. I., Štih, V. and Portugues, R. (2020) “Evidence accumulation during a sensorimotor decision task revealed by whole-brain imaging,” Nature neuroscience. Springer Science and Business Media LLC, 23(1), pp. 85–93.

Gualda, E. J. et al. (2013) “OpenSpinMicroscopy: an open-source integrated microscopy platform,” Nature methods. Springer Science and Business Media LLC, 10(7), pp. 599–600.

Kawashima, T. et al. (2016) “The Serotonergic System Tracks the Outcomes of Actions to Mediate Short-Term Motor Learning,” Cell. Cell Press, 167(4), pp. 933-946.e20.

Keller, P. J., Ahrens, M. B. and Freeman, J. (2015) “Light-sheet imaging for systems neuroscience,” Nature methods. Springer Science and Business Media LLC, 12(1), pp. 27–29.

Khan, B. et al. (2023) “Zebrafish larvae use stimulus intensity and contrast to estimate distance to prey,” Current biology: CB. Elsevier BV, 33(15), pp. 3179-3191.e4.

Kumar, A. et al. (2014) “Dual-view plane illumination microscopy for rapid and spatially isotropic imaging,” Nature protocols, 9(11), pp. 2555–2573.

Lister, J. A. et al. (1999) “nacre encodes a zebrafish microphthalmia-related protein that regulates neural-crest-derived pigment cell fate,” Development, 126(17), pp. 3757–3767.

Patterson, B. W. et al. (2013) “Visually guided gradation of prey capture movements in larval zebrafish,” The journal of experimental biology. The Company of Biologists, 216(Pt 16), pp. 3071–3083.

Petrucco, L. et al. (2023) “Neural dynamics and architecture of the heading direction circuit in zebrafish,” Nature neuroscience, 26(5), pp. 765–773.

Pitrone, P. G. et al. (2013) “OpenSPIM: an open-access light-sheet microscopy platform,” Nature methods. Springer Science and Business Media LLC, 10(7), pp. 598–599.

Semmelhack, J. L. et al. (2014) “A dedicated visual pathway for prey detection in larval zebrafish,” eLife. eLife Sciences Publications Limited, 3, pp. E2391–E2398.

Stelzer, E. H. K. et al. (2021) “Light sheet fluorescence microscopy,” Nature Reviews Methods Primers. Nature Publishing Group, 1(1), pp. 1–25.

Truong, T. V. et al. (2011) “Deep and fast live imaging with two-photon scanned light-sheet microscopy,” Nature methods, 8(9), pp. 757–760.

Vladimirov, N. et al. (2014) “Light-sheet functional imaging in fictively behaving zebrafish,” Nature methods. Nature Research, 11(9), pp. 883–884.

Zylbertal, A. and Bianco, I. H. (2023) “Recurrent network interactions explain tectal response variability and experience-dependent behavior,” eLife, 12. doi: 10.7554/eLife.78381.

